# Simple methods for quantifying super-resolved cortical actin

**DOI:** 10.1101/2021.05.26.445864

**Authors:** Evelyn Garlick, Emma L. Faulkner, Stephen J. Briddon, Steven G. Thomas

**Affiliations:** Institute of Cardiovascular Sciences, College of Medical and Dental Sciences, University of Birmingham, Edgbaston, Birmingham, UK, B15 2TT; Centre of Membrane and Protein and Receptors (COMPARE), University of Birmingham and University of Nottingham, Midlands; Division of Physiology, Pharmacology and Neuroscience, School of Life Sciences, University of Nottingham, Nottingham, UK

**Keywords:** Actin, Super-resolution, SRRF, SIM, Expansion Microscopy, Network analysis

## Abstract

Cortical actin plays a key role in cell movement and division, but has also been implicated in the organisation of cell surface receptors such as G protein-coupled receptors. The actin mesh proximal to the inner membrane forms small fenced regions, or ‘corrals’, in which receptors can be constrained. Quantification of the actin mesh at the nanoscale has largely been attempted in single molecule datasets and electron micrographs. This work describes the development and validation of workflows for analysis of super resolved fixed cortical actin images obtained by both Super Resolved Radial Fluctuations (SRRF) and expansion microscopy (ExM). SRRF analysis was used to show a significant increase in corral area when treating cells with the actin disrupting agent cytochalasin D (increase of 0.31µm^2^ ± 0.04 SEM), and ExM analysis allowed for the quantitation of actin filament densities. Thus this work allows complex actin networks to be quantified from super-resolved images and is amenable to both fixed and live cell imaging.

## Introduction

Cortical actin is a heterogeneous distribution of dense actin filaments in close proximity to the plasma membrane (Clausen et al., 2017), which undergoes constant remodelling (Fritzche et al., 2016) and contributes to cellular structure, migration, and division. Super resolution microscopy studies have indicated that cortical actin can lie between < 10 nm and a maximum of 20 nm from the membrane (Clausen et al., 2017). It is this close association that led to the proposal of the picket fence model by Fujiwara et al. (2002). This model suggests that actin forms small fenced regions, or ‘corrals’, within which membrane proteins can become transiently trapped (Fig. 1). The cytoskeleton is decorated by transmembrane proteins, or ‘pickets’, which have been shown to directly slow local diffusion in the membrane, either by increased packing of lipids around the protein (Sperotto and Mouritsen, 1991), physical steric hindrance (Saxton, 1994) or through increased hydrodynamic interactions (Bussell et al., 1995). There is significant evidence that actin can regulate organisation of certain receptors within the membrane (e.g. serotonin 1a receptor (Shrivastava et al., 2020), α2A-adrenergic receptor (Sungkaworn et al., 2017), and the LYVE-1 receptor (Stanly et al., 2020)) but the precise mechanism by which this occurs has yet to be fully visualised. It would therefore be beneficial to image and analyse these actin corrals at high resolution.

**Figure 1 –.**
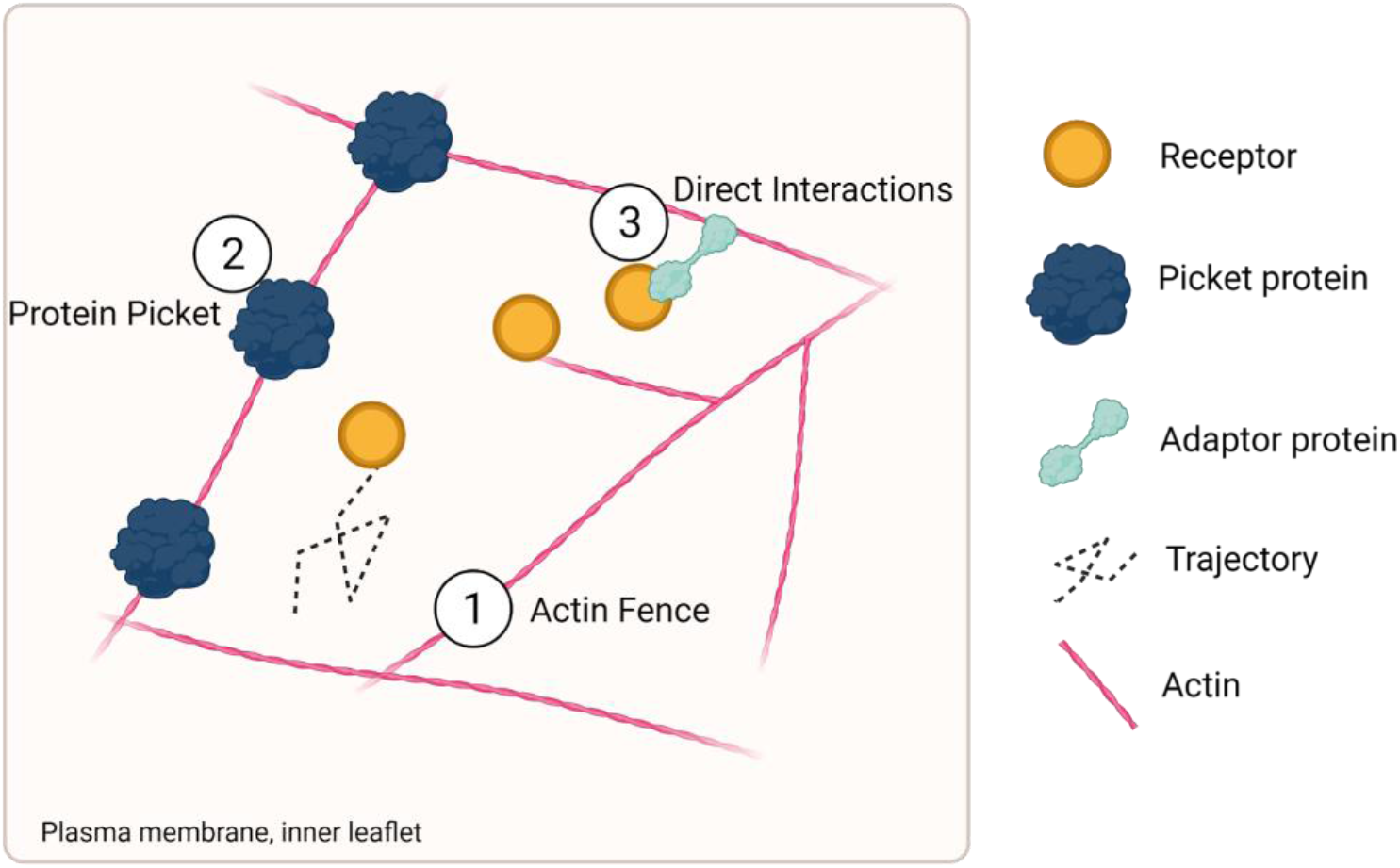
Picket fence model of membrane organisation. Diagram showing proposed nature of the picket fence model on the inner leaflet of the plasma membrane. Potential methods of receptor confinement include 1) direct physical impediment of receptor movement by the actin filaments, 2) alteration to lipid packing around picketing proteins, and 3) direct interaction with actin filaments, with or without adaptor proteins.

Prior to the explosion of super resolution light microscopy techniques, electron microscopy was the best option available to image intricate actin networks at a high resolution. Fujiwara et al. (2016) showed corrals with a median length of 40 nm in PtK2 cells, but 230 nm in NRK cells, suggesting cell specific confinement areas. Electron microscopy does however require complex preparation, which can induce artefacts, as well as having limited options for labelling other molecules of interest. Single molecule localisation microscopy (SMLM) techniques can now resolve structures down to ~20 nm, almost rivalling EM, and therefore capable of resolving individual actin filaments in the dense mesh of the cortical actin (Xu et al., 2012). For example, Photoactivated Localisation microscopy (PALM) imaging of mEOS-Lifeact was used by Sungkaworn et al. (2017) to demonstrate the corralling influence of actin on α2A-adrenergic receptors. Thus far, analysis of super resolved actin filaments largely focuses on single molecule techniques. Peters et al. (2018) developed a method of filament tracing dependent on localisation density, allowing analysis of filament length and branching. Machine learning techniques for point cloud data are also being developed to investigate filamentous structures and their relation to clustered points (Williamson et al., 2020), useful for investigating actin and receptor cluster relationships. Stimulated Emission Depletion (STED) microscopy and Structured Illumination Microscopy (SIM) techniques have a lesser resolving power than SMLM, but still provide at minimum a doubling of standard widefield resolution, and are therefore a powerful tool for actin imaging studies. As a recent example, Stanly et al. (2020) used STED microscopy to image actin corrals in conjunction with the LYVE-1 receptor, reporting corrals with an average size of 100 nm – 1.5 µm, each containing heterogeneously distributed receptor clusters.

As yet, corral structure and function has not been studied in depth by super resolution microscopy in either fixed or live cells. Live actin imaging especially has often relied on widefield and TIRF imaging, which are subject to the resolution limit of light microscopy (~200nm) and therefore unable to resolve finer actin structures. Investigations have therefore largely been on a macro or meso scale, looking at cortical actin intensity, actin flow, or focussing on the large bundled stress fibres. It should not be assumed that all actin can or does contribute in a similar manner to membrane organization - stress fibres often sit a small distance away from the membrane and have very different functions to less bundled cortical actin. However, this does not preclude contribution of larger bundled filaments to membrane protein dynamics (Li et al., 2020).

Analysis of actin networks at a high resolution is a developing field. By focussing on the empty space the actin structure creates, as opposed to the filaments themselves, the work described here sets up analysis workflows that can be used to simply assess the cortical actin mesh in fixed cell super resolved images, as well as quantifying the response of the actin network to disruption.

## Methods

### Cell preparation for SRRF and SIM imaging

A549 cells were plated in #1.5 high tolerance glass bottomed Ibidi dishes and allowed to attach for minimum 18 hours. 1 µM Cytochalasin D was applied for 30 minutes at 37 °C, with equivalent volumes of DMSO as vehicle control, to a final concentration of no more than 0.05%. Cells were fixed for 15 minutes in 4 % PFA in PEM buffer (0.8 M PIPES (pH 6.95), 4 mM EGTA, 1 mM MgSO_4_) that had been pre-warmed to 37 °C, and washed 3x briefly with PBS. Samples were quenched with 50 mM NHCl_4_ for 10 minutes, followed by permeabilization with 0.1 % Triton X-100 for 5 minutes and 3x brief PBS washes. Cells were then labelled with 6.6 nM Alexa Fluor 488 Phalloidin (Invitrogen) for one hour, with 3x brief PBS washes prior to imaging. Imaging was performed in PBS.

Samples were imaged using the Nikon N-SIM-S system (Ti-2 stand, Cairn TwinCam with 2 x Hamamatsu Flash 4 sCMOS cameras, Perfect Focus, Nikon laser bed 488, 561 & 647 nm excitation lasers), Nikon 100x 1.49 NA TIRF Apo oil objective or Nikon 100x 1.35 NA Silicone oil objective, for expansion gels). SIM data was reconstructed using NIS-Elements (v. 5.21.03) slice reconstruction. Representative reconstructed SIM data was assessed by way of the FIJI plugin SIMCheck (Ball et al., 2015). SRRF images were reconstructed from 100 frame bursts of TIRF images, taken at optimal exposure for each sample, as structure rather than intensity was being assessed. For each repeat, a minimum of 10 cells were imaged.

### Expansion Microscopy

Protein retention expansion microscopy was performed according to the technique published by Tillberg et al. (2016). Cells were seeded as above, and fixed with 4 % PFA + 0.1% glutaraldehyde in PEM, prewarmed to 37 °C. Fixation was followed by 3 x PBS washes, quenching of free aldehydes with 50 mM NHCl_4_ for 10 minutes, 3 x PBS washes, and a 5 minute extraction with 0.1% Triton X-100. Where immunofluorescence labelling was necessary, samples were then blocked with a 0.1% Bovine Serum Albumin (BSA)/ 2% goat serum blocking buffer for 1 hour before incubation with the primary antibody, mouse monoclonal anti-α-Tubulin (Sigma-Aldrich), diluted to 1 µg mL^-1^ in blocking buffer, again for 1 hour. After 3 x PBS washes, incubation with the secondary antibody (goat anti-mouse Alexa Fluor 488, A11001, Invitrogen) proceeded for a further hour.

After 3 x PBS washes, all slips were then anchored with 0.1 mg mL^-1^ Acryloyl-X, SE (6-((acryloyl)amino)hexanoic acid, succinimidyl ester (abbreviated as AcX) at room temperature overnight. Slips were washed once in PBS and labelled with Actin ExM (Chrometra) at 1 unit/coverslip for 1 hour. Anchoring with AcX prior to Actin ExM labelling was essential to ensure undistorted retention of the Actin ExM probe post expansion. Gelation proceeded immediately after 3 brief PBS washes.

To make the gel, monomer solution (1x PBS, 2 M NaCl, 8.625 % (w/w) sodium acrylate (SA), 2.5 % (w/w) acrylamide (AA), 0.15 % (w/w) N,N’-methylenebisacrylamide (BIS)) was prepared and aliquoted, storing at −20 °C until use. Concentrated stocks (10 % w/w) of tetramethylethylenediamine (TEMED) accelerator and ammonium persulfate (APS) initiator were added to the monomer solution up to 0.2 % (w/w) each. APS was added last and all solutions were kept on ice to prevent premature gelation. After thorough vortexing of the gel mixture, droplets were pipetted onto parafilm covered glass slides placed in a humid chamber. Coverslips were quickly inverted onto the droplets and gelation was allowed to proceed at room temperature for approximately one minute, before moving to incubate at 37 °C in the dark for 2 hours. It is essential that the humid chamber remains moist during gelation in order to reduce the risk of the gel ripping from the coverslip prematurely in later manipulations. Once gelation was complete, gelled coverslips were carefully removed from the parafilm with flat tweezers and placed in a 6 well plate. Digestion buffer (50 mM Tris (pH 8), 1 mM EDTA, 0.5 % Triton X-100, 0.8 M guanidine HCl) was supplemented with Proteinase K (New England Biolabs), diluted to 8 units mL^-1^. Gelled coverslips were submerged in this digestion buffer overnight at room temperature. Following digestion, gels were transferred to excess deionised water and incubated. Water was replaced until expansion plateaued at an approximate diameter of 5.4cm for the full gel.

For SIM and widefield imaging, high precision glass bottomed Ibidi dishes were fully coated with poly-L-lysine (Sigma-Aldrich) and dried on a hotplate at 95°C. Once fully expanded, a portion of gel was cut with a rectangular coverglass to fit the dish. Excess water was wicked from the gel with tissue to ensure adherence to the coverslip and reduce movement of the gel. The gel was gently pressed once placed on the coverslip to remove bubbles and facilitate firm attachment. A drop of water was added to the top of the mounted gel to minimise shrinkage. Samples were imaged on the Nikon N-SIM-S system as described above.

### Image analysis

Image analysis was performed using Fiji (Schindelin et al., 2012) and Matlab (2019b, v 4.0). Custom written Matlab scripts used for meshwork simulation and line plot analysis are available (https://github.com/biologevie/actin-analysis). A workflow for Fiji analysis is shown in the results section below. SRRF data was assessed with NanoJ-SQUIRREL (Culley et al., 2018a)

### Statistics

All graphs were made and statistical analyses performed using GraphPad Prism 8. Differences between paired groups were assessed using Students T-Test.

## Results

### Generation of super-resolved images of cortical actin networks

SRRF, or Super Resolved Radial Fluctuations (Gustafsson et al., 2016), is a technique that can be used to generate super resolved images from multiple standard widefield, TIRF, or confocal frames. SRRF images of phalloidin stained actin networks in fixed A549 cells were obtained as described in the methods (Fig 2a & b). Fourier Ring Correlation (FRC) measurements show a significant increase in resolution over standard TIRF images (Fig. 2c), as previously reported (Gusstafson et al., 2016, Culley et al., 2018b). Use of NanoJ-SQUIRREL (Culley et al., 2018a) to assess SRRF images indicated robust and accurate reconstruction, routinely reporting Resolution Scaled Pearsons (RSP) correlation coefficient values of >0.95 (Fig. 2d), indicating good reconstruction of SRRF images. Visualisation of error as a heat map indicated issues largely around the thicker filaments (Fig. 2e & f).

**Figure 2 -.**
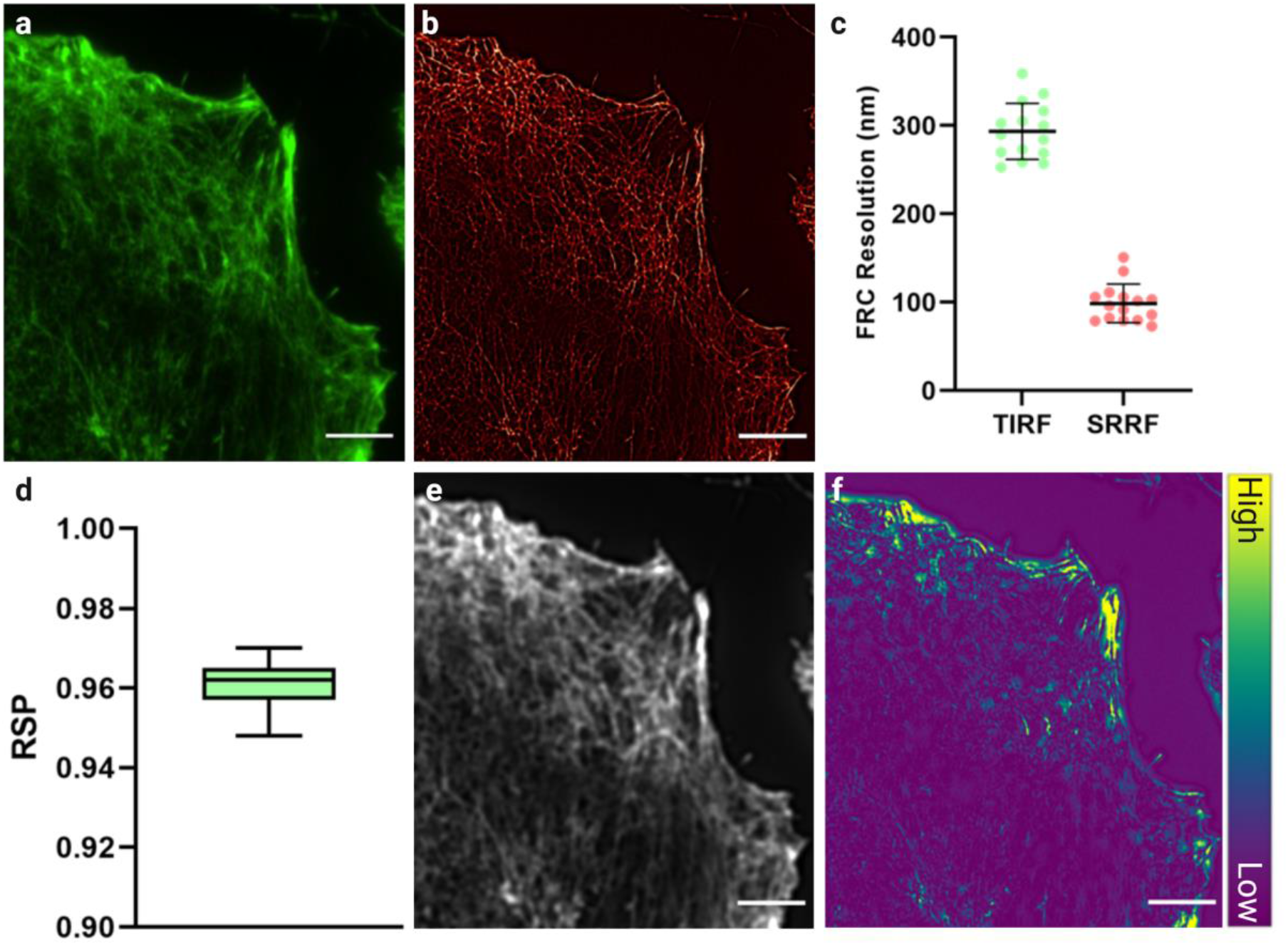
SRRF reconstructions show minimal error compared to TIRF images. a) Representative TIRF image of actin in A549 cells labelled with phalloidin Alexa 488. b) SRRF reconstruction from 100 frames of TIRF data represented in a). c) Mean Fourier ring correlation (FRC) resolution ± standard deviation for the TIRF and SRRF images (n = 15 images from 3 independent experiments). d) Plot showing mean resolution scaled Pearson’s correlation coefficient ± standard deviation for 15 images over 3 independent experiments, as calculated with NanoJ-SQUIRREL. 1 is total correlation and −1 is total anticorrelation. e) Convolved image from b) generated by Nano-J SQUIRREL to calculate error. f) Error map, assessing e) vs a). Indicating artefacts in reconstruction occur in areas of thicker and more dense filaments.

### Quantification of cortical actin networks

In order to quantify the network, a simple method for actin network analysis was tested (Fig. 3a). In FIJI, SRRF images were cropped to an ROI of 10µm^2^, as centrally in the cell of interest as possible (Fig. 3b). The image was then manually thresholded using Otsu’s method. The threshold was used to generate a binary mask of the network, which was followed by erosion (Fig. 3c & d). A classic watershed segmentation was applied (Fig. 3e), and the resulting regions (‘particles’) (Fig. 3e & f) analysed for a range of descriptors, including area and perimeter. To validate accuracy of the analysis, corrals in the original image were measured manually and compared to the thresholded image. This revealed good correlation in terms of the location of corrals identified (Fig. 3f & g), while providing a more consistent assessment of filament delineation. This results in larger estimates of corral area (Fig. 3h - j). This thresholding approach enables greater degree of consistency between images, and a much higher throughput than via manual assessment.

**Figure 3 -.**
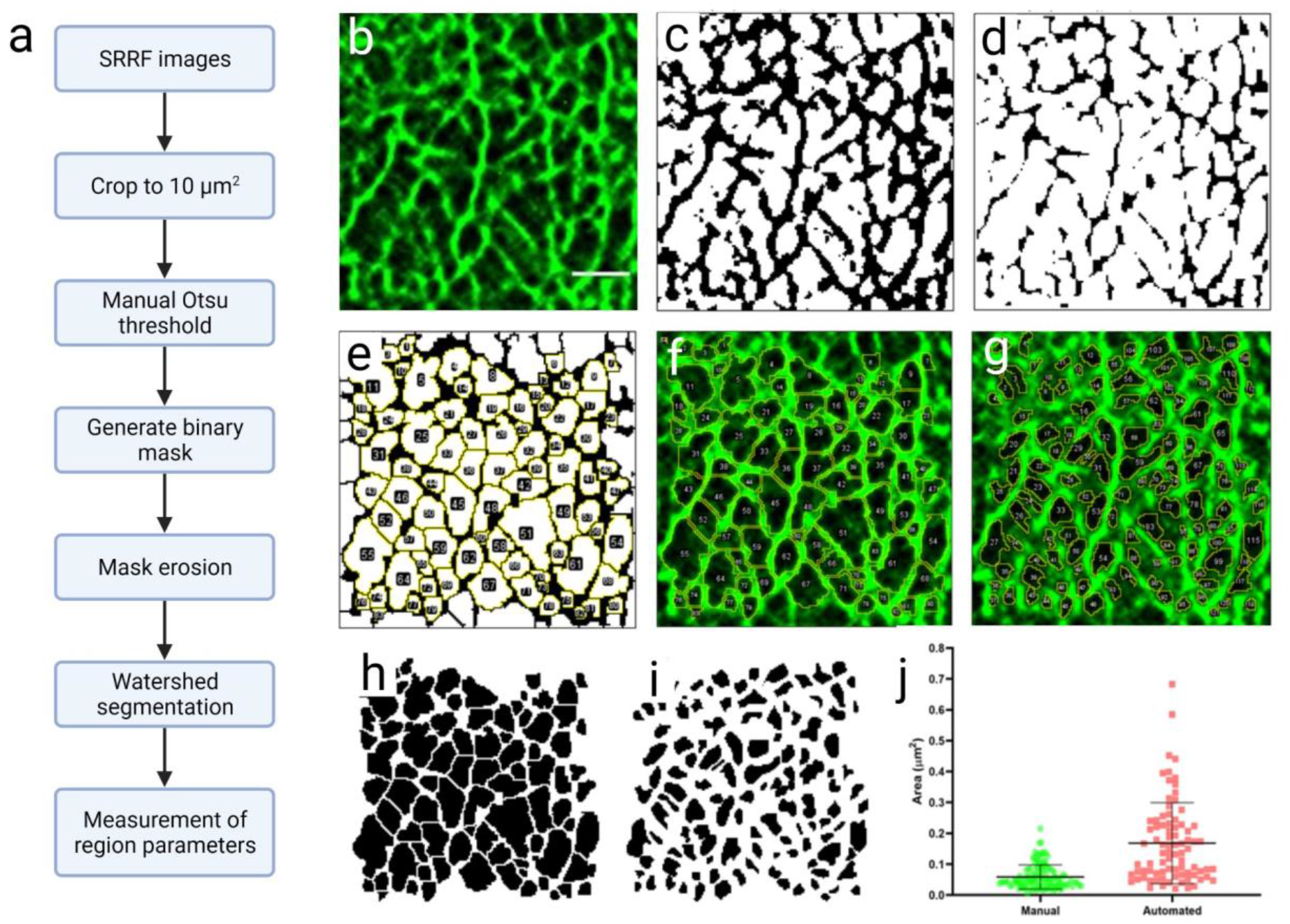
Development of corral analysis workflow. a) Image analysis flow chart, detailing each step taken as illustrated in the following images. b) Cropped SRRF image generated from 100 frames of TIRF data. c) Manually selected Otsu threshold to exclude background pixels. d) Erosion applied to c. e) Watershed segmentation and corral identification (yellow indicates borders of each individually identified region, each numbered). f) Overlay of identified regions on the original cropped SRRF image, compared with g) - manually identified regions hand drawn on the raw SRRF image. h) Mask of regions identified by automated processing, where i) shows masks of regions drawn by hand. j) Comparison of region areas calculated from manual and automated analysis, mean ± standard deviation.

In order to validate the reproducibility of the analysis, we applied it to a ground truth data set. Example actin meshworks were simulated and processed to resemble imaging outputs obtained from our system in terms of pixel size and noise. Image simulation was performed in MATLAB 2019b. Filaments were simulated through random generation of start and end points and each individual filament was given a daughter filament, branching at a 70 degree angle (to mimic Arp2/3 nucleated daughter filaments common in cortical actin networks (Mullins et al., 1998)) (Fig. 4a). Within this code, filament number is user definable, and here was kept at 25 filaments per image, with one daughter filament each. Lines were dilated to more closely resemble the 7nm nature of individual actin filaments and pixels were binned to sizes appropriate for our system and cameras (Fig. 4b) A Gaussian convolution, based on the PSF estimated from our optics, was applied (Fig. 4c). Poisson and Gaussian noise were also applied to give a general approximation of read and shot noise (Fig. 4d), and the image smoothed to give the final simulated image (Fig. 4e). This produced a good representation of our TIRF images of cortical actin networks (Fig. 4f), displaying a similar range of corral sizes within the ground truth data set.

**Figure 4 -.**
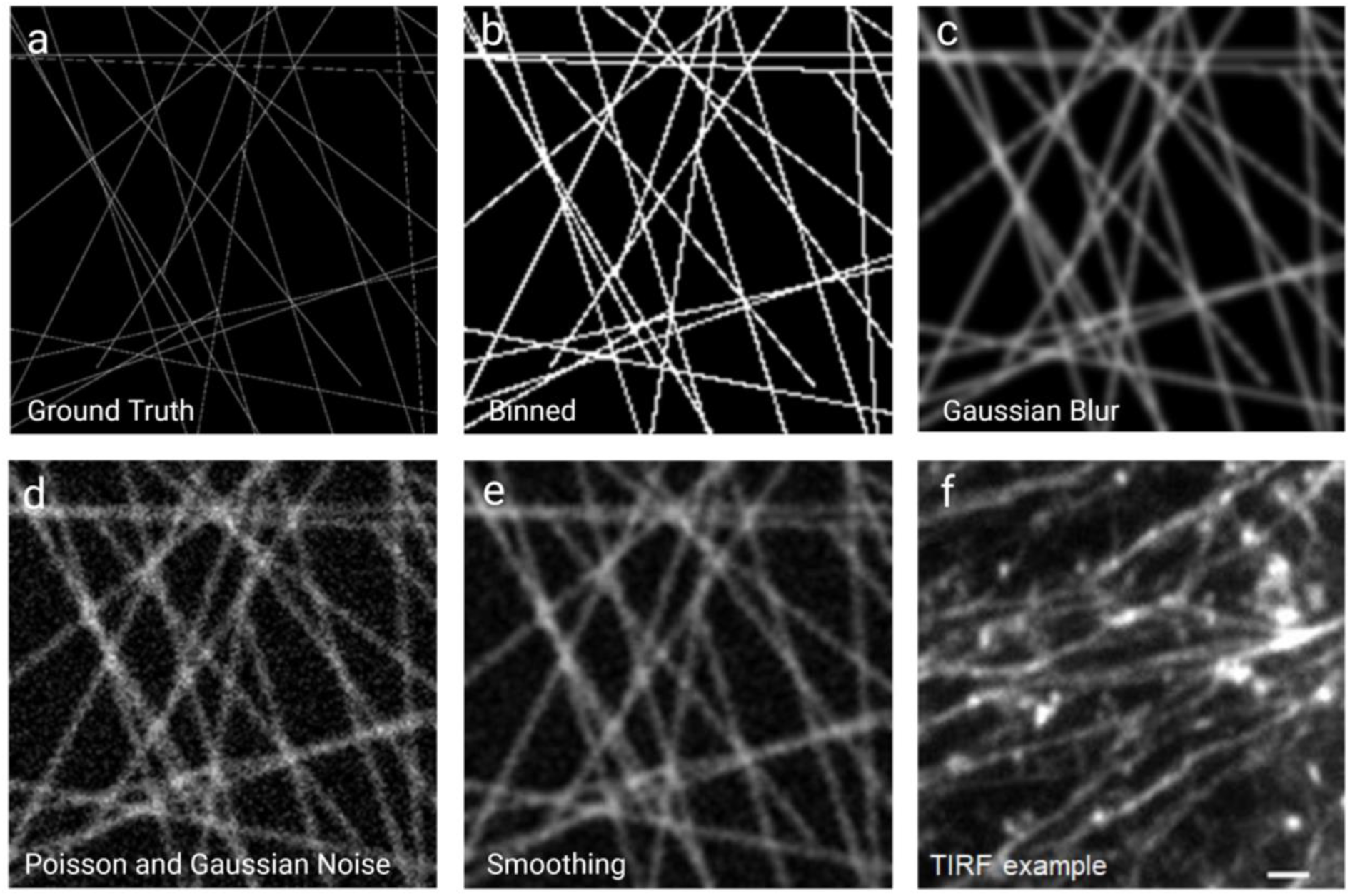
Simulation of meshworks for analysis testing. a) Initial output of 1px wide lines forming a simulated mesh. b) Image a) after dilation of lines to 7px and binning of the image, to match the pixel size of the cameras on our SIM system. c) Convolution with Gaussian blur, as calculated from an estimated PSF based on our optics. d) Application of Poisson and Gaussian noise to mimic noise from the camera. e) Smoothing of d). f) A TIRF image for comparison of the same scale as the simulated data (scale bar = 1 μm).

Analysis of these simulated networks using our thresholding analysis showed good correlation in the identification of the same corrals in the ground truth and simulated data (Fig. 5a & b). Analysis of the area of individual corrals gave similar mean ± SD values in the processed data as for the corrals in the ground truth image (ground truth - 0.51 μm^2^ ± 0.067, processed data 0.49 μm^2^ ± 0.064) (Fig. 5c). The small, but not statistically significant reduction in area in processed images is likely to be a result of the convolution increasing the relative filament thickness, subsequently reducing the corral area. In a similar vein, most incongruities occur with small corrals that become obscured when the PSF is applied. These small corrals can be considered to be below the resolution limit of the images and can be accounted for by filtering corrals under such a size.

**Figure 5 -.**
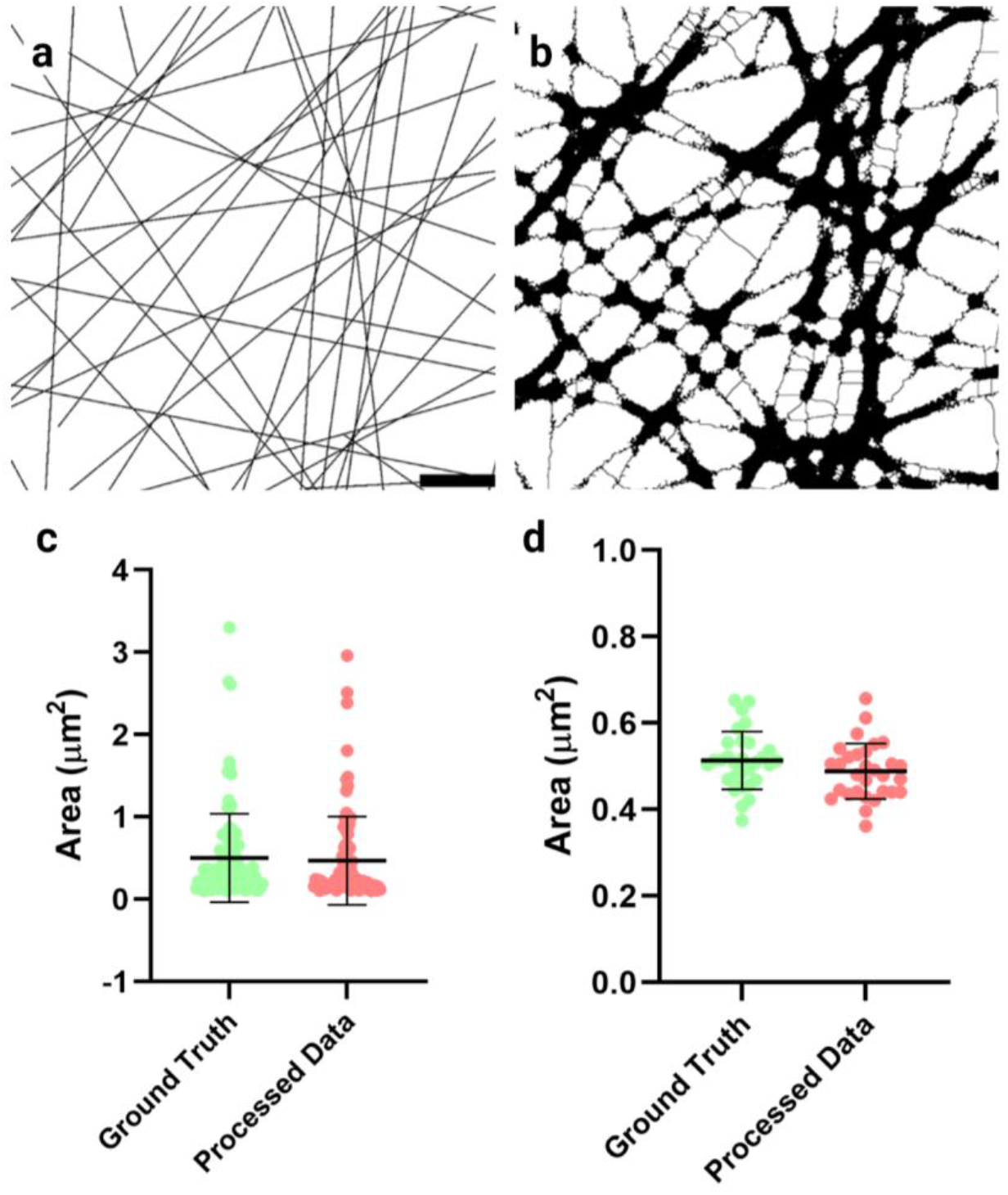
Comparison of ground truth and convolved meshworks shows similar average region areas. a) Ground truth simulated meshwork (scale bar 1.5 μm), shown in b) after convolution and application of our meshwork analysis. c) Areas of regions identified from images a) and b) allowing direct comparison of the meshwork analysis against the ground truth. d) Comparison of mean area ± standard deviation of identified corrals from over 30 repeats of network simulation against the ground truth.

### Cytochalasin D treatment and analysis

To assess the effect of disruption of actin polymerisation on actin corrals and further validate the analysis technique, cells were treated with 1 µM cytochalasin D - a potent inhibitor of actin filament polymerisation prior to fixation, staining and imaging. Representative images of cells ± cytochalasin D are shown in figure 6 (Fig. 6a & b). Cytochalasin D treatment showed clear disruption of normal cortical actin structure. However, to establish if the meshwork analysis could detect changes to the network, a concentration of cytochalasin D was chosen that did not abolish all actin networks (Fig. 6b). Cytochalasin D treatment significantly increased the size of corrals identified by the analysis workflow (Fig. 6c - f). The mean corral area (control - 0.20 μm^2^ ± 0.037, cytochalasin D 0.50 μm^2^ ± 0.19) and perimeter (control - 1.71 μm^2^ ± 0.16, cytochalasin D 2.62 μm^2^ ± 0.48) increased significantly (P<0.0001 for both), while the number of corrals identified in each ROI fell accordingly (control - 386.5 ± 59.39, cytochalasin D 161.5 ± 66.91, P<0.0001).

**Figure 6 -.**
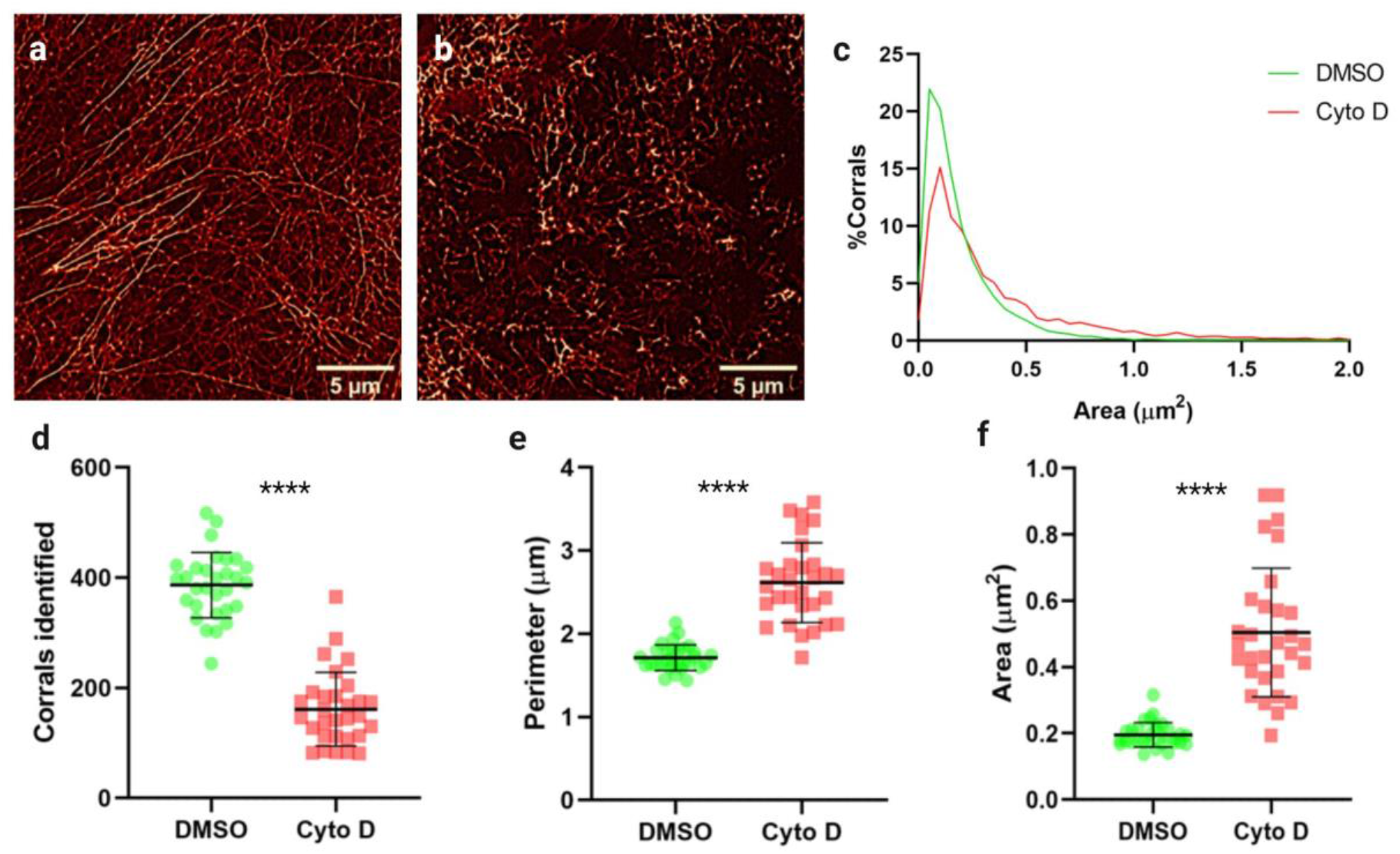
Treatment with cytochalasin D induces significant increase in corral sizes. A) Representative SRRF image of phalloidin labelled actin in DMSO treated cells, and b) cytochalasin D treated cells. c) Histogram of all corral areas for cytochalasin D treated and control cells. d) Number of corrals, e) mean perimeter, and f) mean area of corrals identified in cytochalasin D treated cells vs. DMSO control. n=3, 30 cells per treatment. **** = p<0.0001. All graphs show mean ± standard deviation.

### Expansion microscopy and analysis

While SRRF analysis lends itself to live imaging and is readily adaptable to most imaging setups, resolution is still a limiting factor when considering structures as fine as actin filaments. Given this, we employed expansion microscopy (Tillberg et al., 2016) coupled with SIM imaging to investigate actin networks at resolutions of ~40nm - more comparable to STED’s resolving power, whilst retaining the ability to perform 3D 4-colour labelling for future applications.

In order to visualise actin in expanded cells, the samples were labelled with a phalloidin derivative, Actin ExM. Actin ExM is functionalized with an anchor and therefore compatible with gelation and expansion protocols (Wen et al., 2020). Labelling with this reagent in unexpanded cells shows similar actin structures to those labelled by standard fluorescent phalloidins (Fig. 7). 3D-SIM images of expanded samples clearly show actin structure, with fine meshworks and thicker bundled fibers well labelled (Fig. 8a). 3D reconstructions also highlight the ability of ExM to visualise cortical actin filaments from filaments deeper in the cell (Suppl. Video 1).

**Figure 7 -.**
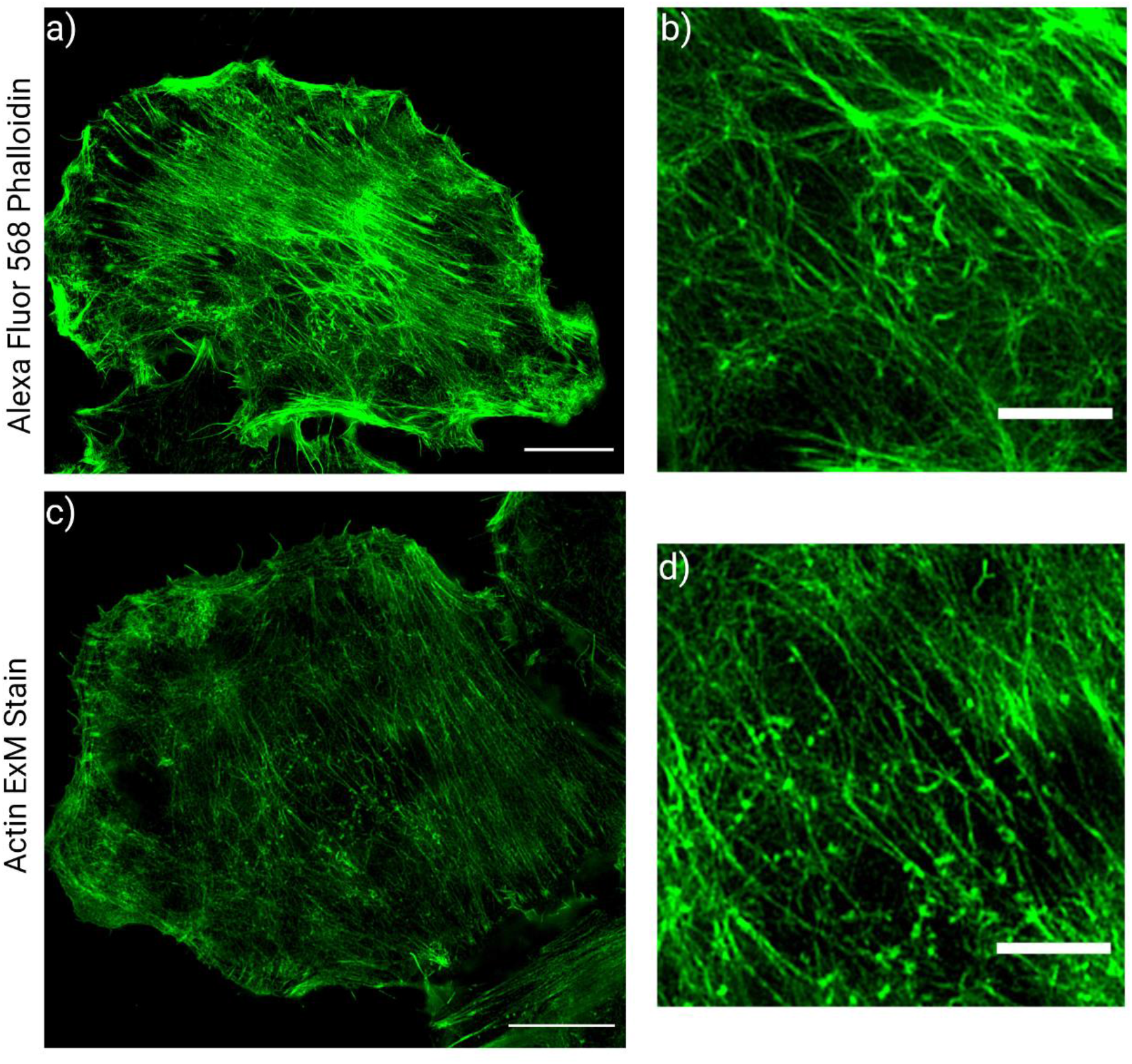
Comparison of Actin ExM labelling reagent and AlexaFluor 568 phalloidin. a) Representative 3D SIM reconstruction of an A549 cell labelled with standard AF 568 phalloidin, with b) showing a zoomed in section of the central region of the cell. c) Representative 3D SIM reconstruction of an unexpanded, un-gelled A549 cell labelled with Actin ExM, with d) showing a zoomed in section of the central region of the cell. (Scale bars: a & c = 10 μm, b & d = 4 μm)

**Figure 8 -.**
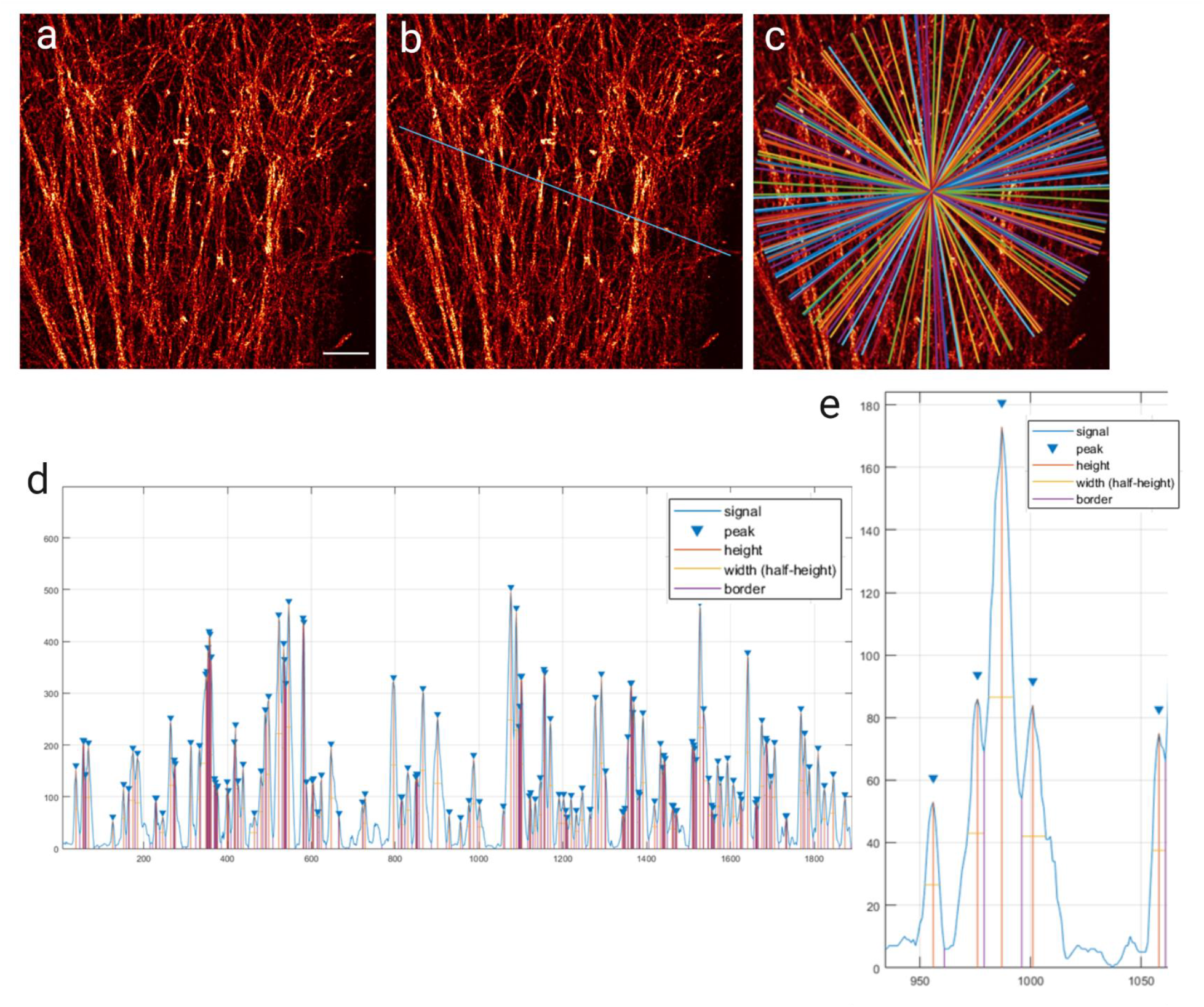
Example of analysis of expanded actin. a) Example of 3D-SIM imaged expanded actin, with a scale bar of 10 μm (uncorrected for expansion factor). b) Randomly generated line across the image for analysis, where intensity along this line is plotted in d). c) Example of the distribution of randomly generated analysed lines when repeated 100 times. d) Normalised intensity line plot of fluorescence intensity (a.u.) vs pixels of line in b, with peaks identified and height and width denoted as in the key. e) Enlarged region of d), showing identification of all distinguishable peaks above a manually set threshold.

Analysis of expanded samples presented a number of challenges. Some discontinuity in filament labelling, as well as the contribution of out of focus light (although minimised by adjustment of SIM reconstruction parameters) made application of the analysis used for SRRF images difficult. As such, density of filaments was approximated by examining fluorescence intensity across ROIs of 3D-SIM reconstructions. Randomly oriented lines across full ROIs were drawn (Fig 8b) and fluorescence intensity graphs generated. These values were normalised and local maxima identified (Fig 8d), with a threshold to eliminate background, but no threshold on peak prominence, allowing resolution of closely adjacent yet still separate filaments (Fig 8e). Where peaks were not clearly delineated, a straight line was drawn to separate each peak, and width at half prominence calculated from these (Fig. 8d). Repeating this operation hundreds of times over a single image (Fig 8c) and comparing the numbers of peaks calculated for each ROI served as a way to compare filament density across the image.

To demonstrate the power of this analysis, images of A549 cells stained for microtubules and actin and imaged via ExM-SIM (Fig 9a & b) were subject to filament density analysis. Visually the two networks are seen to comprise different densities of filaments. Quantifying the number of peaks identified showed that the actin network consisted of a higher number of filaments than the microtubule network (Fig 9c & d). This was confirmed by calculating the distance between peaks (Fig. 9 e & f), again showing that the actin networks are more dense than the microtubule ones. This data shows that filament density analysis can determine subtle differences in the organisation of filaments in a cell.

**Figure 9 -.**
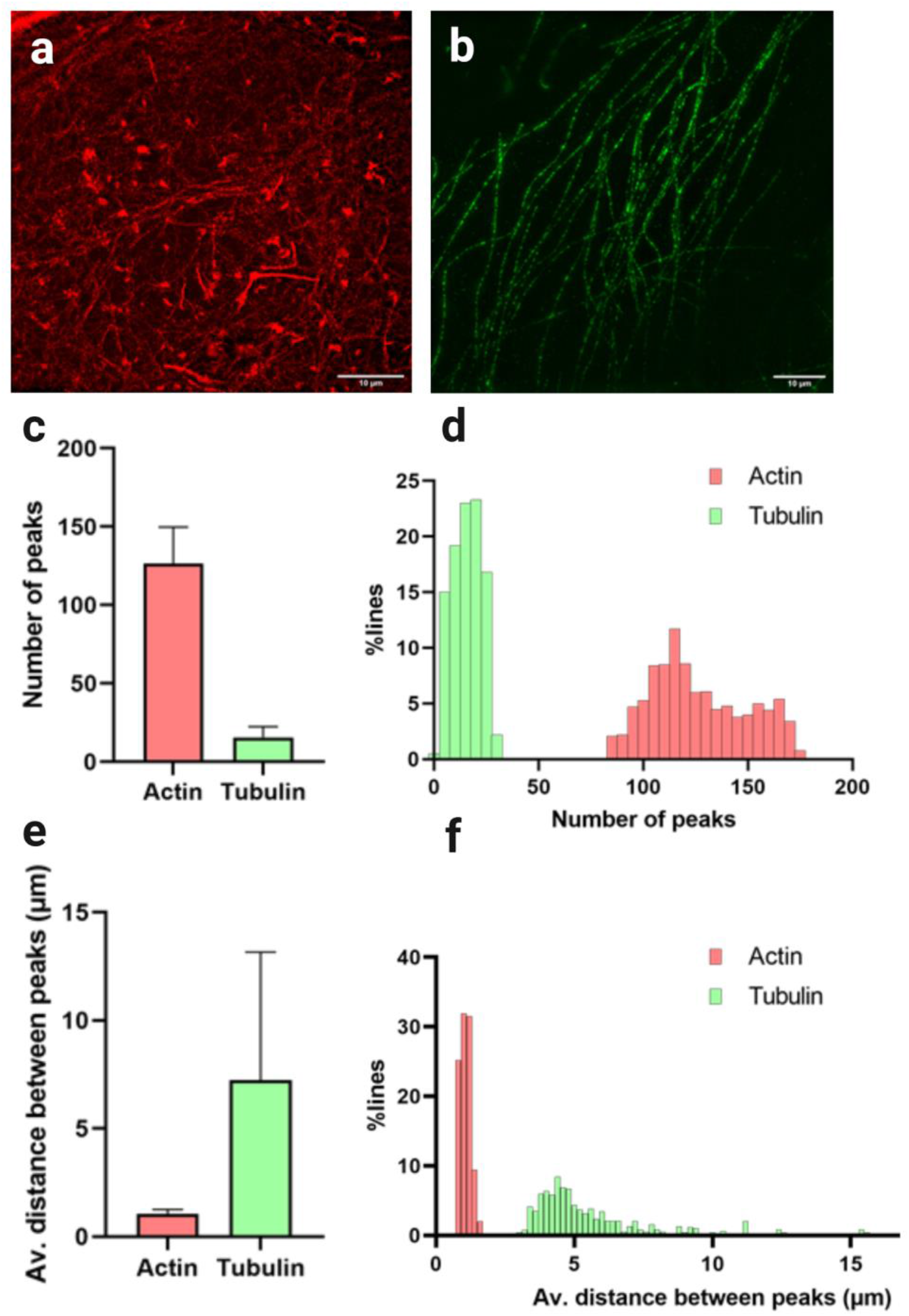
ExM filament analysis distinguishes between small and large meshworks. a) Example 3D SIM image of expanded actin and b) of expanded tubulin (both scale bars 10 μm). c) Mean number of peaks ± standard deviation. identified over 100 repeats of the analysis for each image, with d) showing a histogram of this dataset. e) Mean distance between identified peaks ± standard deviation over 100 repeats of the analysis for each image, with the histogram shown in f. Lengths in e and f are not corrected for expansion factor.

## Discussion

In this study we have developed and tested the application of two workflows for quantifying cortical actin organisation in super-resolved microscopy images. It contrasts other actin filament analysis methods in that, rather than skeletonising the filaments and calculating filament length, filament branching, branching angle, etc., our SRRF analysis assesses the shapes and sizes of the meshwork.

Accurate quantitation of the actin meshwork depends on the generation of reliable, quality super-resolved images. SRRF imaging performs well on actin, giving clear and reproducible structures that correlate well with the TIRF raw data and as quantified by SQUIRREL analysis. However, false sharpening and the collapsing of adjacent objects is a common issue in SR reconstructions, including in common single molecule localisation microscopy reconstruction algorithms (Marsh et al., 2018). Care should be taken when reconstructing super-resolution images and appropriate quality control checking performed. The application of SQUIRREL (Culley et al, 2018a) here helped to identify regions of possible error in reconstructions. These were mostly observed in brighter, sharper regions of the reconstruction. However, it is important to note that the error maps are only internally relative; RSP values give a more inter-experiment and inter-technique comparable value, and in our experiments RSP values were consistently high.

The meshwork analysis developed here performs well on SRRF data, and on data simulated to represent the binning and pixel size of SRRF data. The comparison of simulated versus ground truth data also highlighted that the analysis can underestimate some areas by over segmenting as well as missing smaller regions that sit below the resolution limit. This is, however, an issue inherent to any resolution limited image, and by filtering out very small corrals (i.e. those below the resolution limit) from further analysis. With this caveat in mind, the analysis performed well, identifying similar regions, which were also similar in size, between our simulated and ground truth data. To further test the analysis, and to see if it could detect subtle changes in the actin network, we treated cells with the actin disruptor cytochalasin D. From our analysis, the effect of cytochalasin D on the samples was as predicted – a visible disruption of normal cortical actin structure, with the persistence of some, especially thicker, filaments. Cytochalasin D acts by binding the barbed end of actin filaments, inhibiting polymerisation and dissociation from this end. Typically, cytochalasin D induces the formation of denser focal accumulations of actin, likely by interrupting normal anchoring of filaments by capping proteins (Cooper, 1987). As such, normal membrane associations should be severely disrupted, as indicated by our data. The accurate representation of cytochalasin D effects by our analysis workflow therefore strengthens justification for broader application and to assess subtle changes in cortical actin network organisation in response to inhibitors, stimulation, etc..

Skeletonisation analysis methods work well for continuous, well separated filaments, such as microtubules, intermediate filaments, or even thicker actin stress fibers (see Zhang et al. (2017) for implementation of their technique SFEX to quantify actin stress fibers in TIRF images). When considering super resolved fine actin meshworks, however, discontinuity and reduced fluorescence intensity is an inherent issue. This can lead to artefacts like mismatching of filament segments and artificial removal of sparser filaments. By focussing on the gaps the actin leaves rather than trying to extract information from the filaments themselves, artefacts introduced in the thresholding steps can be somewhat mitigated. A recent preprint describes an algorithm called FiNTA that performs more advanced and accurate filament tracing than typical skeletonisation (Flormann et al., 2021), but this technique is reported to perform best on filaments of uniform thickness, an area where our analysis appears more robust.

The simple technique we describe should be easily applicable to other super resolution techniques, such as SIM and STED, but better options are available for SMLM. As a rule of thumb, where analysis can be applied directly to the point cloud data generated in single molecule techniques rather than images reconstructed from this data, it should be. This retains the maximum information and resolution gained by using these techniques. Peters et al. (2018) developed an algorithm to perform filament tracing from the spatial point patterns generated in SMLM imaging of actin, allowing extraction of information directly from the point cloud rather than the reconstructed image. Identification and differentiation of fibrous and clustered structures in SMLM data is also possible using Williamson et al.’s (2020) machine learning based analysis approach.

Expansion microscopy offers a far superior resolution to SRRF, albeit only in fixed cell contexts. Underutilised in the study of actin, expansion microscopy is shown here to faithfully preserve both bundles and fine actin structure, with resolution almost comparable to SMLM when used in conjunction with SIM (Wang et al, 2018). Where expansion could be arguably an improvement on single molecule techniques is the potential for simple and rapid multiplexing of labels. The nature of ExM as, essentially, a modified standard immunofluorescence technique, means that imaging four spectrally distinct labels in one sample is relatively easy to achieve. While techniques like DNA-PAINT (Jungmann et al., 2010) are more applicable than dSTORM or PALM for multi-colour imaging, this comes at a significant time cost - a single image can take hours to acquire. In addition, as expansion microscopy sample preparation optically clears the sample, making samples more amenable to super-resolution 3D imaging, expansion is a strong candidate for investigating complex actin structures in 3D in cells. Recently, Park et al. (2020) used anti-fluorophore antibodies to image actin labelled with standard Alexa-488 conjugated phalloidin, which, while a different approach to the one taken here, indicates that techniques for actin imaging in expanded samples are growing. Whilst filament tracing routines have been applied to microtubules post expansion (Flormann et al., 2021), the relative density and continuity of fine actin filaments continues to restrict application. The method we describe here uses fluorescence intensity fluctuations as a proxy for filament density and allows these complex filament networks to be quickly quantified.

In conclusion, we demonstrate two simple analysis methods for quantifying cortical actin networks in super-resolved microscopy images. These methods allow for quick, reproducible quantitation of actin corral number and size in SRRF or SIM images, and for quantitation of filament density in ExM images. These methods are amenable to batch processing of large data sets, and to adaptation for live cell analysis of actin dynamics.

## Acknowledgements

We thank the British Heart Foundation (IG/18/2/33544; NH/18/3/33913) and COMPARE for funding. The authors would like to acknowledge the Imaging Suite at the University of Birmingham for support of imaging experiments. Imaging facilities used in this project were funded by the University of Birmingham, COMPARE and the BHF.

## Author Contributions

These contributions follow the Contributor Roles Taxonomy guidelines: **https://casrai.org/credit/**

Conceptualization: E.G., S.J.B., S.G.T

Funding acquisition: S.J.B.,S.G.T.

Investigation: E.G., E.L.F

Methodology: E.G.

Software: E.G.

Supervision: S.J.B., S.G.T.

Resources: E.G., E.L.F

Visualization: E.G., S.G.T.;

Writing – original draft: E.G., S.G.T.;

Writing – review & editing: E.G., E.L.F., S.J.B., S.G.T.

All authors had the opportunity to read and comment on the final manuscript.

## Notes

### Competing Interest Statement

The authors have declared no competing interest.

