## Supplemental information for "Simple methods for quantifying super-resolved cortical actin"

### Supplementary Information

Supplementary video 1

Movie showing a portion of an Expanded A549 cell labelled with Actin ExM and imaged using 3D-SIM. Depth colour scale applied allowing actin filaments at different depths to be identified.
